# The basement membrane regulates the cellular localization and the cytoplasmic interactome of Yes-Associated Protein (YAP) in mammary epithelial cells

**DOI:** 10.1101/2023.08.23.554465

**Authors:** Antonio Carlos Manucci, Ana Paula Zen Petisco Fiore, Giovani Luiz Genesi, Alexandre Bruni-Cardoso

**Author notes:** Co-authors details: Antonio Carlos Manucci, current address: Department of Biochemistry, Institute of Chemistry, University of São Paulo, São Paulo, Brazil; Ana Paula Zen Petisco Fiore, current address: Center of Genomic and System Biology, Department of Biology, New York University, New York, NY, USA; Giovani Luiz Genesi, current address: Department of Neurology, Perelman School of Medicine, University of Pennsylvania, Philadelphia, PA, USA.

## Abstract

The Hippo pathway, a signaling cascade involved in the regulation of organ size and several other processes, acts as a conduit between ECM cues and cellular responses. We asked whether the basement membrane (BM), a specialized ECM component known to induce quiescence and differentiation in mammary epithelial cells, would regulate the localization, activity and interactome of YAP, a Hippo pathway effector. To address this question, we used a broad range of experimental approaches, including 2D and 3D cultures of both mouse and human mammary epithelial cells, as well as the developing mouse mammary gland. In contrast to malignant cells, non-tumoral cells cultured with a reconstituted BM (rBM) displayed higher concentrations of YAP in the cytoplasm. Incidentally, when in the nucleus of rBM-treated cells, YAP resided preferentially at the nuclear periphery. In agreement with our cell culture experiments, YAP exhibited cytoplasmic predominance in ductal cells of developing mammary epithelia, where a denser BM is found. Conversely, terminal end bud (TEB) cells with a thinner BM displayed higher nucleus-to-cytoplasm ratios of YAP. Bioinformatic analysis revealed that genes regulated by YAP were overrepresented in the transcriptomes of microdissected TEBs. Consistently, mouse epithelial cells exposed to the rBM expressed lower levels of YAP-regulated genes, although the protein level of YAP and Hippo components were slightly altered by the treatment. Mass spectrometry analysis identified a differential set of proteins interacting with YAP in cytoplasmic fractions of mouse epithelial cells in the absence or presence of rBM. In untreated cells, YAP interactants were enriched in processes related to ubiquitin-mediated proteolysis, whereas in cells exposed to rBM YAP interactants were mainly key proteins related to amino acid, amino sugar and carbohydrate metabolism. Collectively, our multifaceted approach unraveled that the BM induces YAP translocation or retention in the cytoplasm of non-tumoral epithelial cells and that in the cytoplasm YAP seems to undertake novel functions in metabolic pathways.

## Introduction

Chemical and mechanical signals provided by the extracellular (ECM) regulate a milieu of cellular processes. Reciprocally, cells remodel the ECM by differentially secreting and/or degrading ECM molecules and by altering the ECM topology (Cerqueira et al., 2022; Walma & Yamada, 2020). This complex molecular interplay, involving bidirectional communication between the extracellular matrix (ECM) and cells, plays a critical role in tissue development, homeostasis, and proper function, and is disrupted in diseases such as cancer (Fiore et al., 2018; Walma & Yamada, 2020). Signals from the ECM modify the tension and organization of the cytoskeleton and modulate intracellular pathways triggered by soluble factors (Hastings et al., 2019, 2019; Kadry & Calderwood, 2020).

The ECM is chemically and topographically heterogeneous among organs and within the same organ (Hastings et al., 2019). For example, the basement membrane (BM), a specialized ECM compartment composed mainly of laminins, collagen type IV, nidogen and perlecan, surrounds epithelia and other tissues providing anchorage and cues that influence essential molecular processes determining cellular outputs (Inman et al., 2015). In the mammary gland, the convergence of signals from sex hormones, other soluble systemic and organ-intrinsic factors, and cues from the BM activate cellular quiescence and differentiation towards milk production (Inman et al., 2015). Nevertheless, our understanding of the mechanisms through which signals from the BM are transmitted to the cell nucleus regulating the genome remains incomplete.

A candidate to bridge the epithelial BM to cell nucleus communication is the Hippo pathway. In mammals, the main members of the pathway are the serine-threonine kinases MST1/2 and LATS1/2, the scaffold proteins SAV1 and MOB1A/B and the effectors YAP and TAZ (Gumbiner & Kim, 2014). Canonically, when Hippo is active, MST1/2 phosphorylates SAV1 assembling a complex with it; the MST1/2-SAV1 complex then phosphorylates MOB1A/B, which in turn recruits LATS1/2 so that it is also phosphorylated by MST1/2 (Dasgupta & McCollum, 2019). Activated LATS1/2 phosphorylates YAP and/or TAZ (Lei et al., 2008), forcing their retention in the cytoplasmic (Kim & Jho, 2018) . Once sequestered in the cytoplasm, YAP/TAZ can be ubiquitinated by the E3-ligase SCF^βTrCP^ complex and degraded via proteasome 26S (Kim & Jho, 2018). When the Hippo pathway is inactive, unphosphorylated YAP/TAZ can translocate to the nucleus and act mainly as co-activators of the TEAD family of transcription factors (TEAD1/2/3/4), stimulating the expression of genes related to proliferation, survival and stemness (Zhao, Li, Lei, et al., 2010). Nevertheless, there is robust evidence supporting that in some mechanical contexts, YAP/TAZ localization is independent of Hippo (Dupont et al., 2011).

We used a combination of 3D cell culture, fluorescence microscopy, proteomics, and bioinformatics to investigate whether the BM would regulate the localization, activity and interactome of YAP in mammary epithelial cells. The nucleus-to-cytoplasm ratio of YAP was lower at the ducts of developing mammary epithelial cells, which are abundantly surrounded by the BM. In non-tumoral cells treated with a reconstituted BM (rBM), YAP was predominantly localized at the cytoplasm, but when in the nucleus, it dwelled preferentially close to the nuclear envelope. Genes regulated by YAP were overrepresented in the terminal end buds (TEB) of the developing mammary gland and in 3D tumor-like structures formed by human breast cancer cells. Furthermore, mouse epithelial cells exposed to the rBM expressed lower levels of YAP-regulated genes, albeit the level of YAP and Hippo components were not altered by the treatment. We identified a differential set of proteins interacting with YAP in cytoplasmic fractions of mouse epithelial cells in the absence or presence of rBM. The YAP interactants found in non-treated cells are mainly involved with ubiquitin-mediated proteolysis, whereas in cells exposed to rBM proteins interacting with YAP were mainly key regulators of cellular metabolism. Altogether, our work revealed that signals from the BM alter the localization and interactome of YAP and provides important clues for novel cytoplasmic functions of YAP in physiologically relevant settings.

## Results

We sought to investigate whether the basement membrane (BM) would alter the cellular localization and interactome of YAP in mammary epithelial cells. Consistent to what has been shown for different types of epithelial cells in subconfluent monolayer cultures (Das et al., 2016), YAP staining was largely localized in the nuclei of EpH4 cells (a mouse mammary non-malignant epithelial cell line) (Fig. 1A). However, in a tridimensional (3D) model composed of a reconstituted BM (3D-rBM) in which cells assemble into differentiated epithelia resembling mammary acini, YAP was predominantly cytoplasmic (Fig. 1B). This localization pattern was also reproduced in S1 cells, a non-malignant human breast cell line of the HMT3522 progression series which in 3D-rBM also form acinus-like structures (Fig. 1C) (Rizki et al., 2008). In T4-2 cells, a malignant cell line also derived from the HMT3522 series which in 3D-rBM develops into tumor-like structures instead of acini, YAP was homogeneously distributed in the cells, with no preferential nuclear or cytoplasmic localization (Fig.1C-D). T4-2 cells present aberrant proliferation and polarity signaling pathways, but they can be phenotypically reverted upon inhibition of a single disrupted pathway (Fiore et al., 2017; Furuta et al., 2018; Wang et al., 2002; Weaver et al., 1997). T4-2 reverted (T4-2R) cells are sensitive to BM signaling and arrange in growth-arrested epithelial structures similar to non-malignant cells. Accordingly, YAP localization in T4-2-reverted (T4-2R) structures was analogous to what we found for S1 acini (Fig. 1C-D). Taken together, these cell culture findings indicate that in non-tumoral contexts mammary epithelia respond to the growth-repressive and differentiating cues from the BM promoting the reduction of nuclear YAP, thus likely decreasing the expression of genes regulated by this transcriptional co-activator.

**Figure 1.**
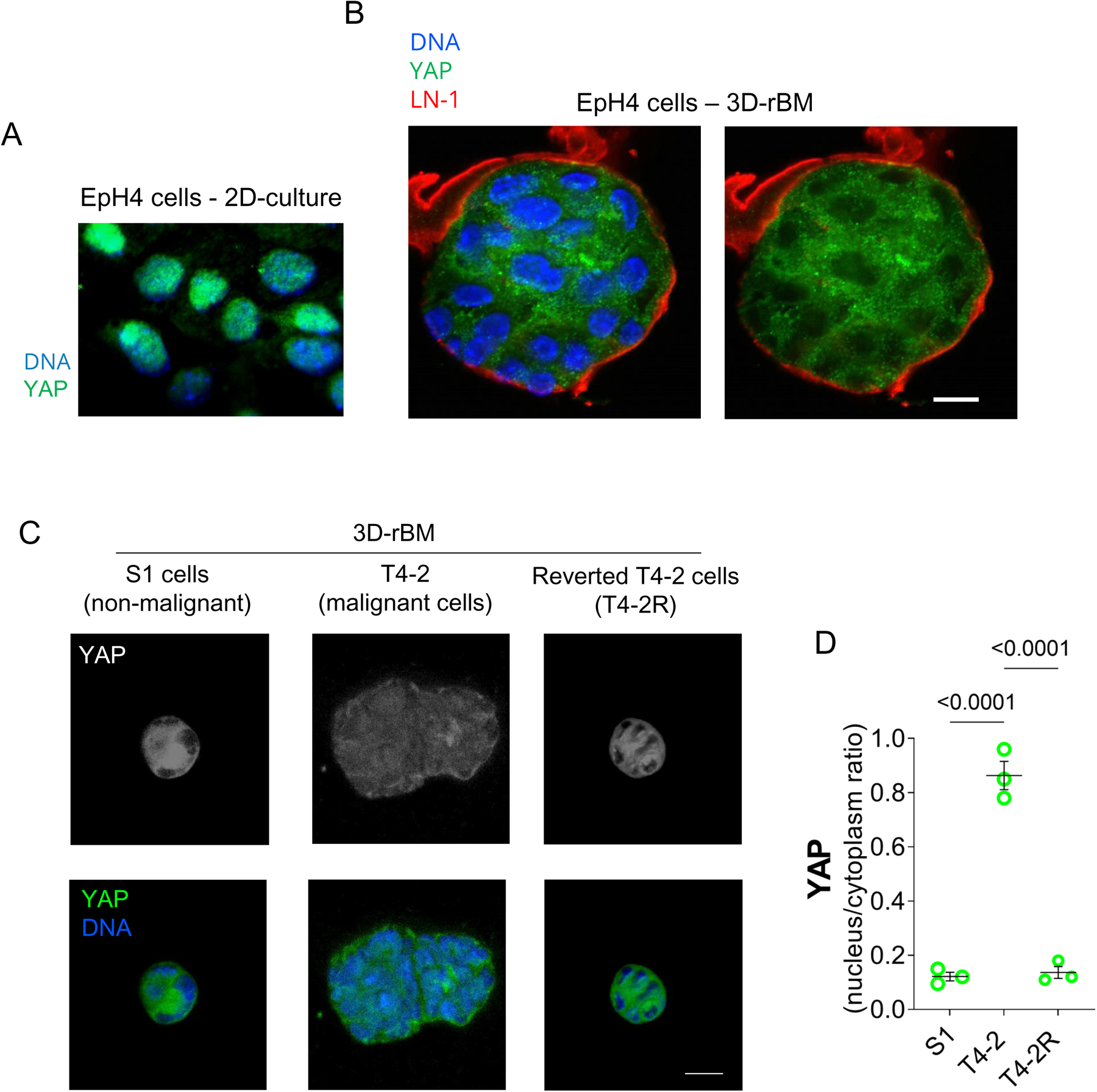
A 3D-rBM induces cytoplasmic retention of YAP. A) Immunofluorescence confocal image of a subconfluent EpH4 monolayer (2D) culture. YAP (green) is predominantly localized in the nucleus (DNA stained with DAPI; blue). B) Cryosection of a 3D EpH4 cell cluster cultured in rBM. In this condition, YAP (green) is more cytoplasmic. Laminin-111 (LN-111; red) staining indicates the assembly of BM encasing the 3D spheroid. DNA stained with DAPI (blue). Scale bar = 10 μm. C) Immunofluorescence for YAP of the human breast epithelial cells S1 (nonmalignant), T4-2 (malignant), and phenotypically reverted T4-2-cells (T4-2R) cultured in a 3D-rBM for 5 days. As opposed to nonmalignant and phenotypically reverted cells, T4-2-cell structures display a higher nucleus to cytoplasm ratio of YAP. D) Quantification of YAP nucleus-to-cytoplasm ratio in the human cells grown in 3D. Data in bar graph are represented as mean +/-SEM. The data were submitted to ANOVA followed by Tukey posthoc test. ** = p<0.01. N= 3. Scale bars = 20 μm.

We asked whether YAP localization *in vivo* would corroborate cell-culture findings. The developing mammary gland shows dynamic remodeling of the BM and consists in an excellent model to study the molecular regulations that the BM exerts on various cellular processes. In accordance to previous work (Fiore et al., 2017; Inman et al., 2015; Polyak & Kalluri, 2010), the terminal end buds (TEBs) - the growing tips of the developing mammary epithelial tissue containing progenitor and proliferative cells - presented a thinner BM, as revealed by a fainter laminin-111 staining (Fig. 2A). On the other hand, the relatively quiescent and differentiated ducts of the same epithelial structures displayed a thicker BM (Fig. 2A). YAP was predominantly nuclear at the TEBs, whereas in the ducts YAP nucleus-to-cytoplasm ratio was lower (Fig. 2A-B). Consistent to what we found for cells grown surrounded by an rBM, a lower nucleus-to-cytoplasm ratio of YAP was associated with the thick BM found in the developing mammary ducts (Fig. 2B).

**Figure 2.**
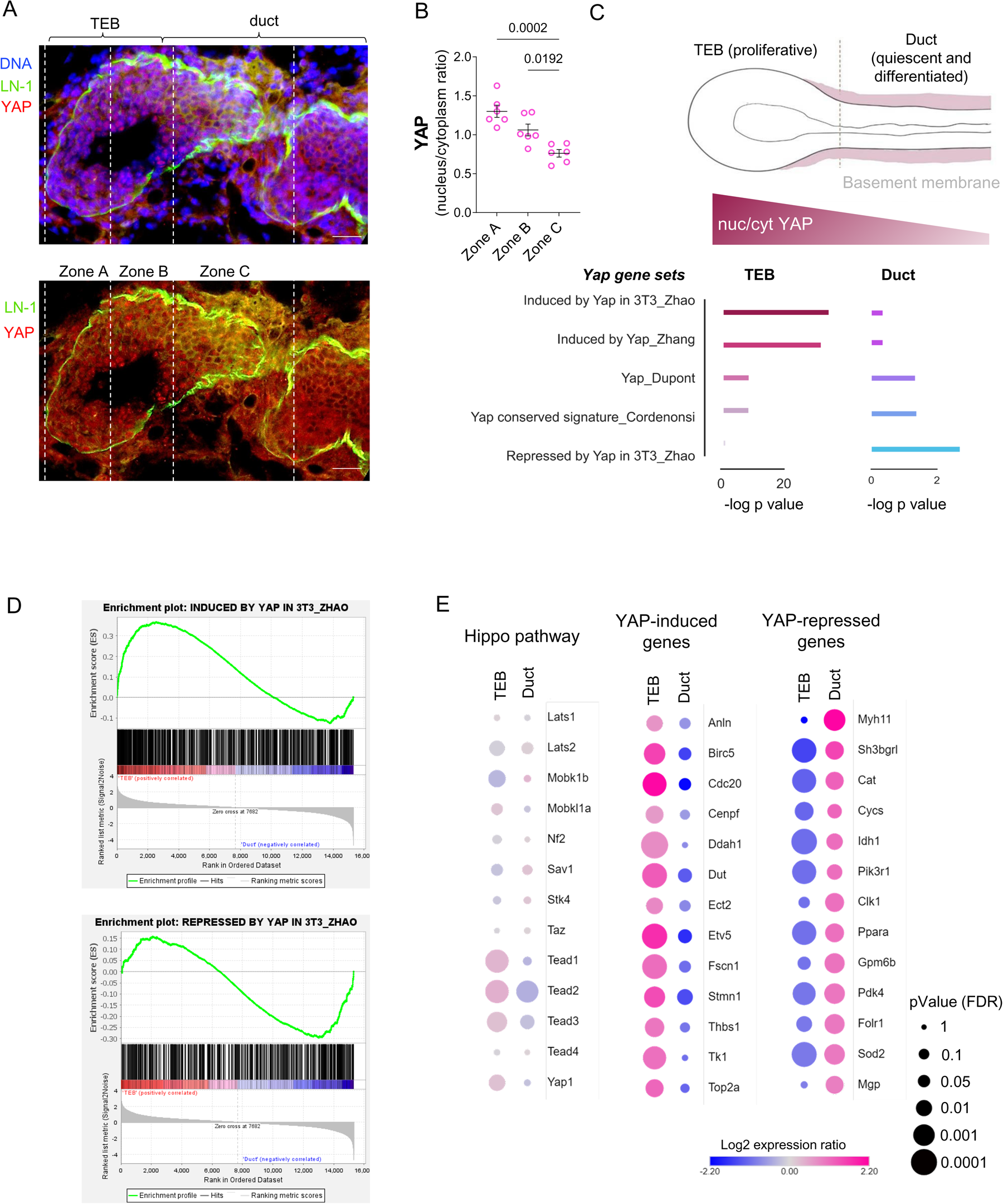
The mammary terminal end buds (TEB) present a higher nucleus-to-cytoplasm ratio of YAP and overrepresentation/enrichment of YAP gene signatures. A) A developing mammary gland epithelium (Balb/C 5-week year old) stained for Yap (red), laminin-111 (LN-1; green) and DNA (DAPI). LN-1 surrounds the epithelia and shows an intensity staining gradient fashion being stronger at the duct microenvironments. B) Quantification of Yap nucleus to cytoplasm ratio in 3 different zones of mammary epithelia (6 TEBs from 3 animals; 2 TEBs/animal). The 3 different zones were determined using the beginning of TEBs as a start point. Zones A and B are 40 µm and zone C is 80 µm in width. We analyzed 20-55 cells per zone. Data in the plot shows measurements of each TEB (dots) and mean +/-SEM. The data were submitted to ANOVA followed by Tukey posthoc test. * = p< 0.05; **= p<0.01. Scale bar = 20 µm. C) Overrepresentation (ORA) analysis performed with Enrichr for YAP regulated genes in the gene expression profiles of TEB and duct (Kouros-Mehr & Werb, 2006). YAP regulated gene sets were retrieved from the literature (see material and methods). C-top) Schematic of a developing mammary epithelium highlighting a thicker BM surrounding duct cells and a higher YAP nucleus to cytoplasm ratio in cells within TEB. C-bottom) The bar plot shows ORA for TEB (lef) and duct (right). Data are shown as -log of the adjusted p-value (FDR). Yap induced gene sets are only overrepresented in TEB, while YAP-repressed genes are overrepresented only in duct. D) Gene set enrichment analysis (GSEA) of Yap regulated genes in the gene expression profiles of TEB and duct. Representative GSEA enrichment plot for Yap induced gene sets and an enrichment plot of Yap repressed genes. E) Changes in mRNA levels of Hippo pathway genes, Yap-induced genes and Yap-repressed genes in TEB and duct. The Hippo pathway genes were curated from literature. Yap-induced genes are the 13 genes from the YAP_Dupont gene set overrepresented in TEB. Yap-repressed genes are the 13 genes with the highest expression and overrepresented in duct from the repressed by Yap in 3T3_Zhao gene set. The data in the dot plot are shown as log_2_ of TEB ratio (= M(TEB) – M(duct)), converted to fold change, and Duct ratio (= M(duct) – M(TEB), converted to fold change). See the color code bar illustrates the fold-change magnitude. The dot sizes in the plot reflect -log_10_ of adjusted p values (FDR).

To assess if YAP localization in the developing mammary gland correlated with differential expression of YAP-regulated genes, we assessed a public dataset generated by the Zena Werb laboratory (Kouros-Mehr & Werb, 2006) in which number 4 mammary glands of 5-week-old were microdissected into terminal end bud (TEB), mature duct, and epithelium-free distal stroma components for microarray transcriptomics analysis (see material and methods of the original work (Kouros-Mehr & Werb, 2006). We considered as differentially expressed genes within TEB or duct expression ratios higher than 1.5-fold change and adjusted p-value < 0.05 (Benjamini– Hochberg correction, also known as False Discovery Ratio (FDR)). Using these cutoffs, we found 604 differentially expressed genes in TEB and 508 genes in duct. An overrepresentation analysis (ORA) with the bioinformatics tool Enrichr revealed that sets of genes whose expression are induced by YAP (YAP-induced gene sets) were significantly overrepresented in TEB profiles (Fig. 2C, Suppl. Table 1a). A gene set enrichment analysis (GSEA) using the whole gene expression list from the Kouros-Mehr and Werb (Kouros-Mehr & Werb, 2006) dataset confirmed that YAP-induced gene sets were indeed enriched in TEB (Fig. 2D and Suppl. Fig. 1A). Consistently, a list of genes whose expression is repressed by YAP (YAP-repressed gene set) was only overrepresented and enriched in duct gene expression profile (Fig. 2C-D and Suppl. Fig. 1B). We also interrogated the dataset in respect to the expression of Hippo pathway components (Fig. 2E). For this, we manually curated from the literature a list of genes that are core members of the Hippo pathway. Intriguingly, most Hippo pathway genes were not differentially expressed, and even for genes with an FDR below 0.05 had low effect sizes, with no genes showing TEB or Duct ratios higher than 1.5 or lower than 0.5 (Fig. 2E - left column). Next, we inspected the 13 genes of the YAP_Dupont gene signature found in the TEB gene expression profile and found that they indeed had higher expression in TEB, in sharp contrast to the 13 top genes from the YAP-repressed signature that were more expressed in duct (Fig. 2E – middle and right columns). Taken together, our data revealed that mRNA expression of Hippo pathway genes is not differential in TEB, a niche poor in BM, while both localization of YAP and expression of YAP-regulated genes are sharply distinct TEB and duct microenvironments.

As described above, non-malignant, and phenotypically reverted mammary epithelial cells in 3D-rBM had lower nucleus-to-cytoplasm ratios of YAP, raising the question whether that was due to the rBM signaling itself or to the 3D environment created in the assay. To tackle this issue, we adapted a 3D cell culture assay where cells form clusters on non-adhesive plates and are treated or not with rBM (Fig. 3A and reference (Xu et al., 2009)). The model was validated by assessing traits presented by EpH4 cells grown in conventional 3D-rBM cultures supplemented with lactogenic hormones (Xu et al., 2009). Most clusters assembled in smaller and more spherical tissue structures when exposed to rBM (Fig. 3B-D) and about 60% of the structures presented lumen and formed typical 3D cell culture acini as assessed by the presence of visible lumen (Fig. 3E). Non-treated clusters, on the other hand, showed irregular 3D geometries and rarely displayed lumen (Fig. 3B-E). β-casein, a marker of functional differentiation of mammary epithelial cells, was robustly expressed at the transcriptional and protein level only in EpH4-cell clusters treated with rBM (Fig. 3F-G).

**Figure 3.**
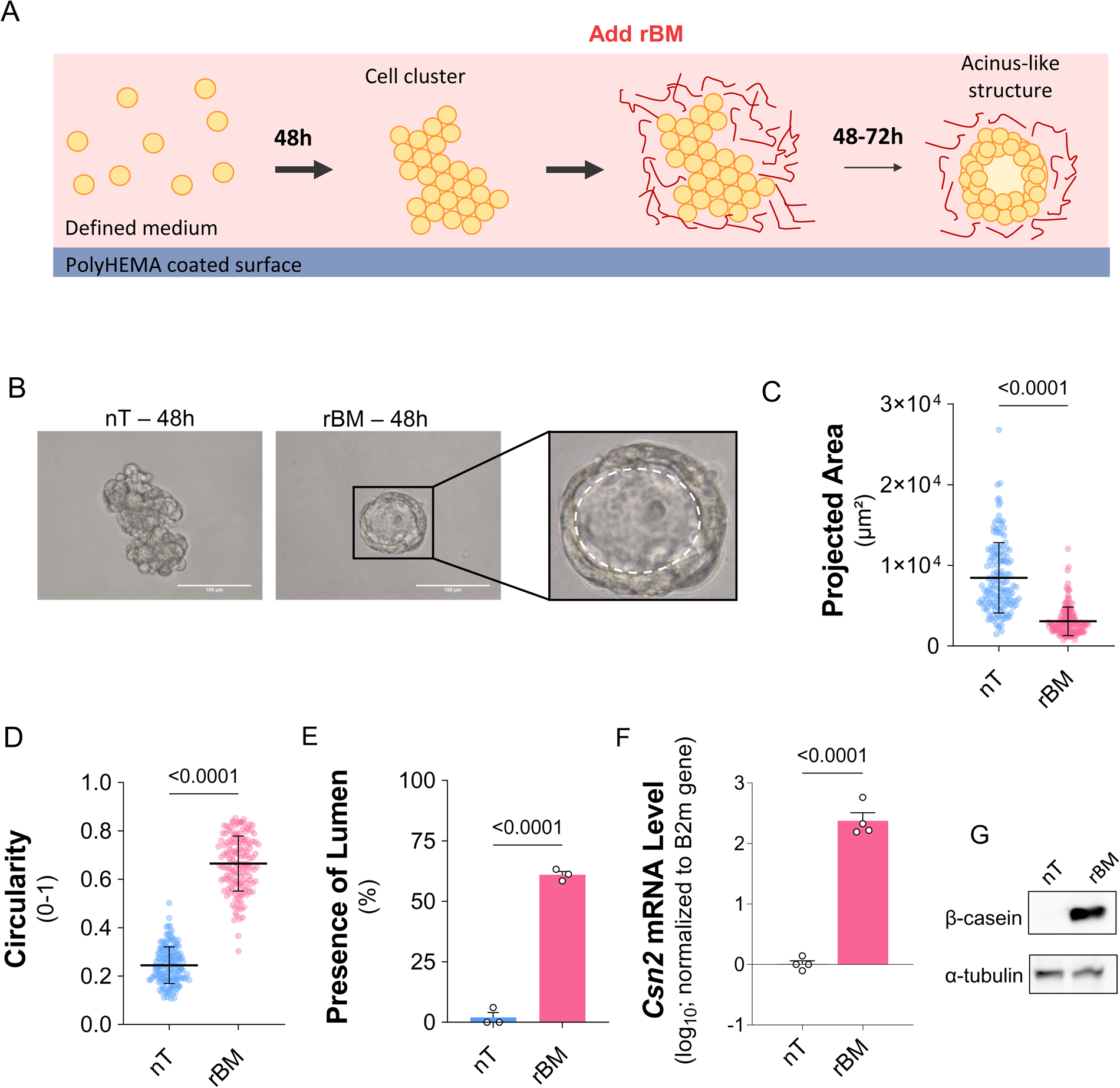
A tridimensional model to probe the role of the BM on YAP subcellular localization. A) Schematic of the tridimensional culture over non-adhesive surfaces. B) Phase-contrast micrographs of non-treated and rBM-treated 3D EpH4 clusters after 48h. Inset: An rBM-treated epithelial cell structure with its lumen highlighted (dashed line). Scale bar = 100 μm. C-E) Morphological analysis of 3D mammary epithelial structures: projected area (C), circularity (D), and presence of lumen (E). F-G) Cell differentiation was assessed by quantification of mRNA by RTqPCR (F) and protein levels by western blotting (G) of β-casein, the main protein component present in milk. The data in the plots are shown as mean +/-SD (C and D) or SEM (E and F). The data were submitted to two-tailed unpaired T-test; N=3.

We found a distinct pattern of YAP distribution when comparing rBM-treated and non-treated EpH4-cell 3D structures. The nucleus-to-cytoplasm ratio of YAP was higher in non-treated cells, confirming that it is not the 3D environment itself that induces changes in YAP localization, but signaling from the BM (Fig. 4A-B). Strikingly, in this model a ring of YAP staining at the nuclear periphery in cells within clusters treated with rBM became evident (Fig. 4C). Double labeling of YAP and lamin-B, a marker for the nuclear lamina, together with tracing profiles of fluorescence intensity unveiled that the YAP “ring” was inside the nucleus, but with little overlap with DAPI-stained DNA (Fig. 4C-D). This indicates that YAP is not only translocated from the nucleus to the cytoplasm in presence of rBM, but also the remaining nuclear YAP is redistributed within the cell nucleus. The mRNA levels of *Mst2,* a core Hippo kinase, *Yap* and crucial YAP-target genes, *Ankrd1* and *Ect2* significantly reduced in 3-rBM cultured cells (Fig. 5A). Although the protein levels of MST2 and YAP were slightly decreased, other key components of Hippo as well as phosphorylation of MOB, LATS and YAP, which are a measure of the pathway activation status, were not changed with the treatment (Fig. 5B-C). These data are consistent with a higher cytoplasmic presence of YAP found in 3D-rBM treated cells and indicate that our 3D cell culture model reproduced the pattern seen for the developing mammary epithelia and that the level of Hippo proteins and their activation are not altered by the BM, albeit YAP localization, and consequently its activity, are strongly controlled by the BM.

**Figure 4.**
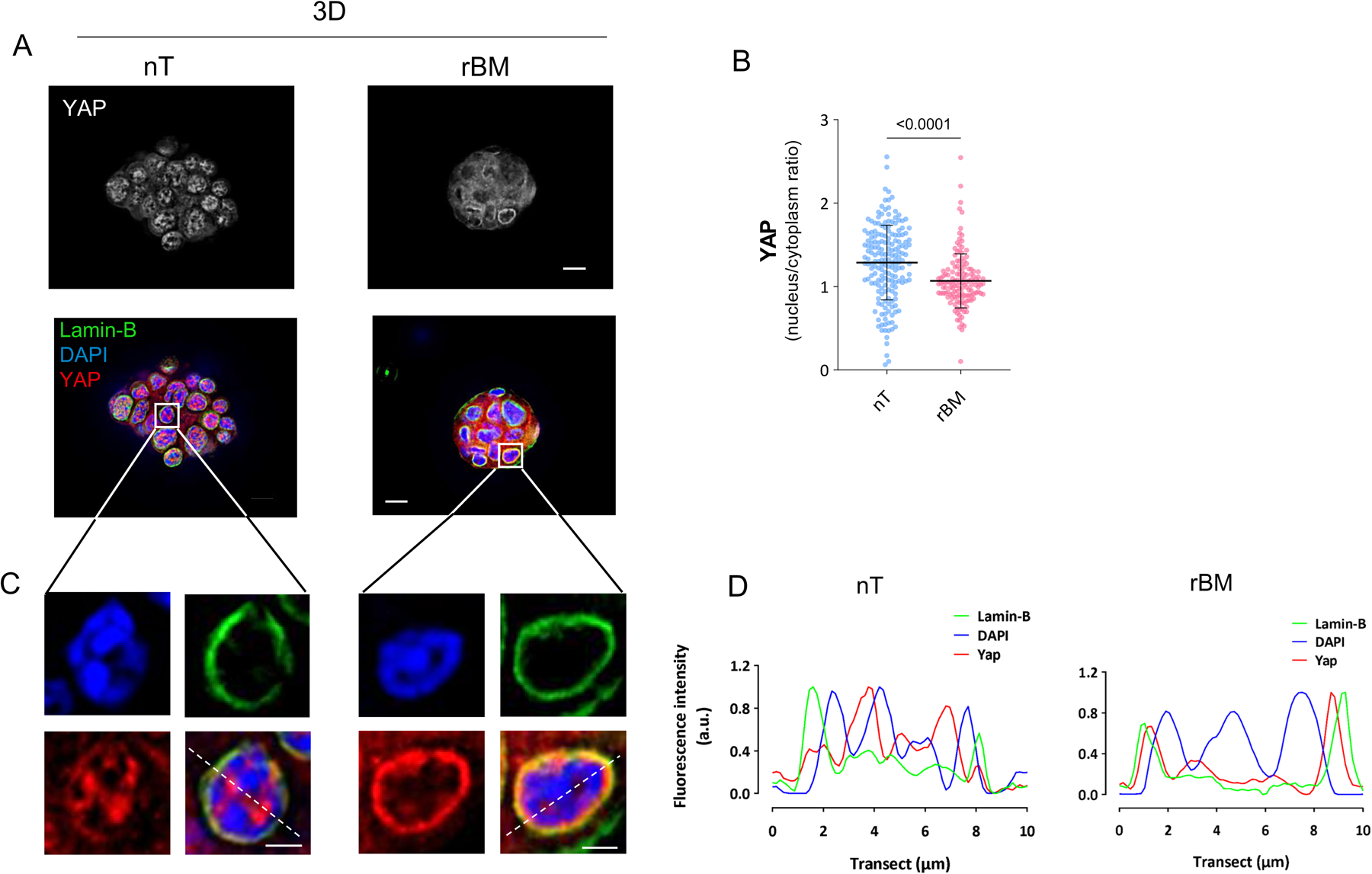
3D-rBM alters YAP distribution in mammary epithelial cells. A) Fluorescence micrographs of 3D EpH4 cell structures cultured for 72 hours in the absence (nT) or presence of an rBM submitted to immunostaining of lamin-B (green) and YAP (gray or red). DNA was stained with DAPI. B) YAP nucleus-to-cytoplasm ratio quantification. C) Insets of the images displayed in A. D) Fluorescence intensity profile for lamin-B, YAP and DAPI staining intensities, based on the white dashed lines overlaid in C. The data in the plots are shown as mean +/-SD. The data were submitted to two-tailed unpaired T-test; N=3.

**Figure 5.**
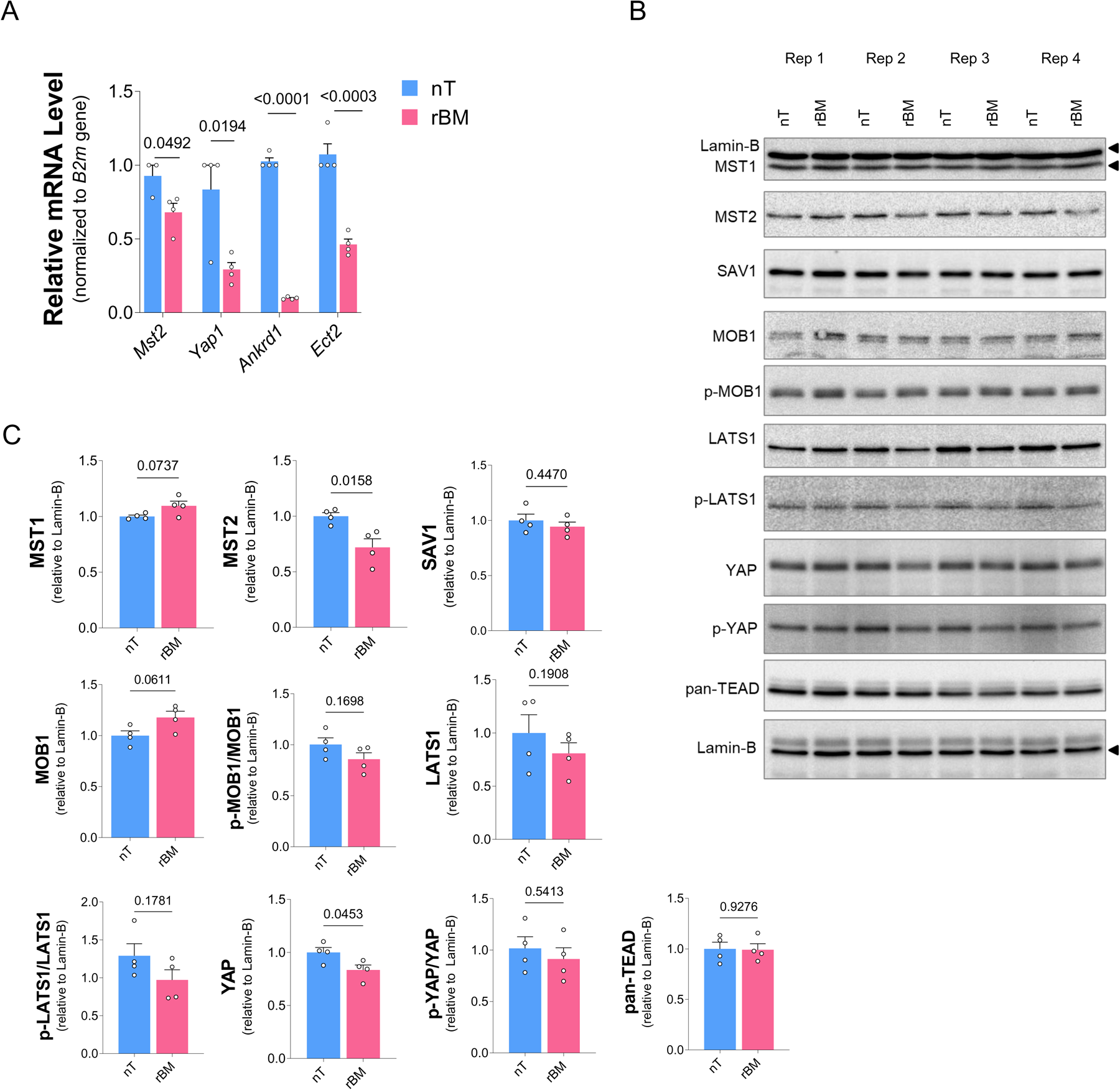
3D-rBM decreases expression of YAP target genes despite inducing subtle changes in protein level and phosphorylation of key components of the Hippo pathway. A) RT-qPCR analysis of *Stk3* (Mst2 encoding gene), *Yap* and *Ankrd1* and *Ect2* (target genes of YAP) gene expression relative to *B2m* expression in 3D EpH4 cell structures cultured in the absence (nT) or presence of rBM. B) Western blots (WB) of Hippo pathway proteins of 3D EpH4 cell structures cultured in the same condition as in B). All four independent replicates (Rep) were run in the same gels and are presented in displayed in the panel. Lamin-B was used as a loading control for the western blots. Note that the band appearing above MST1 (first blot) refers to lamin-B, which was detected before MST1, however not stripped away from the blot membrane. C) Quantification of western blots shown in B). Data in the plots are shown as mean +/-SEM and data were submitted to two-tailed unpaired T-test (p-values shown in the plots); N=3 for the RT-qPCR and =4 for the western blots.

We became intrigued with YAP localization in the cytoplasm of epithelial cells in the duct of the developing mammary gland and in nonmalignant epithelial cells in 3D rBM and asked if YAP would have a biological function in the cytoplasm of epithelial cells. To generate insights regarding this question, we performed cellular fractionation and mapped the interactomes of YAP in the cytoplasmic fraction of EpH4 cells using co-immunoprecipitation of endogenous YAP followed by mass spectrometry (Fig. 6A-B). Because this type of strategy requires a scale that is unfeasible for 3D cell cultures, we used the ECM overlay assay (Fiore et al., 2017; Streuli et al., 1995; Tomasin et al., 2023) in which cells were grown under a defined medium for 48h and then rBM and prolactin were added to the medium (Fig. 6A). In this assay, important features seen in 3D-rBM are reproduced, such as a change from flat to rounded cell morphology, cell cycle arrest and expression of β-casein (Spencer et al., 2011; Streuli et al., 1995; Tomasin et al., 2023), and importantly, the nucleus to cytoplasm ratio of YAP is decreased (Suppl. Figure 2).

**Figure 6.**
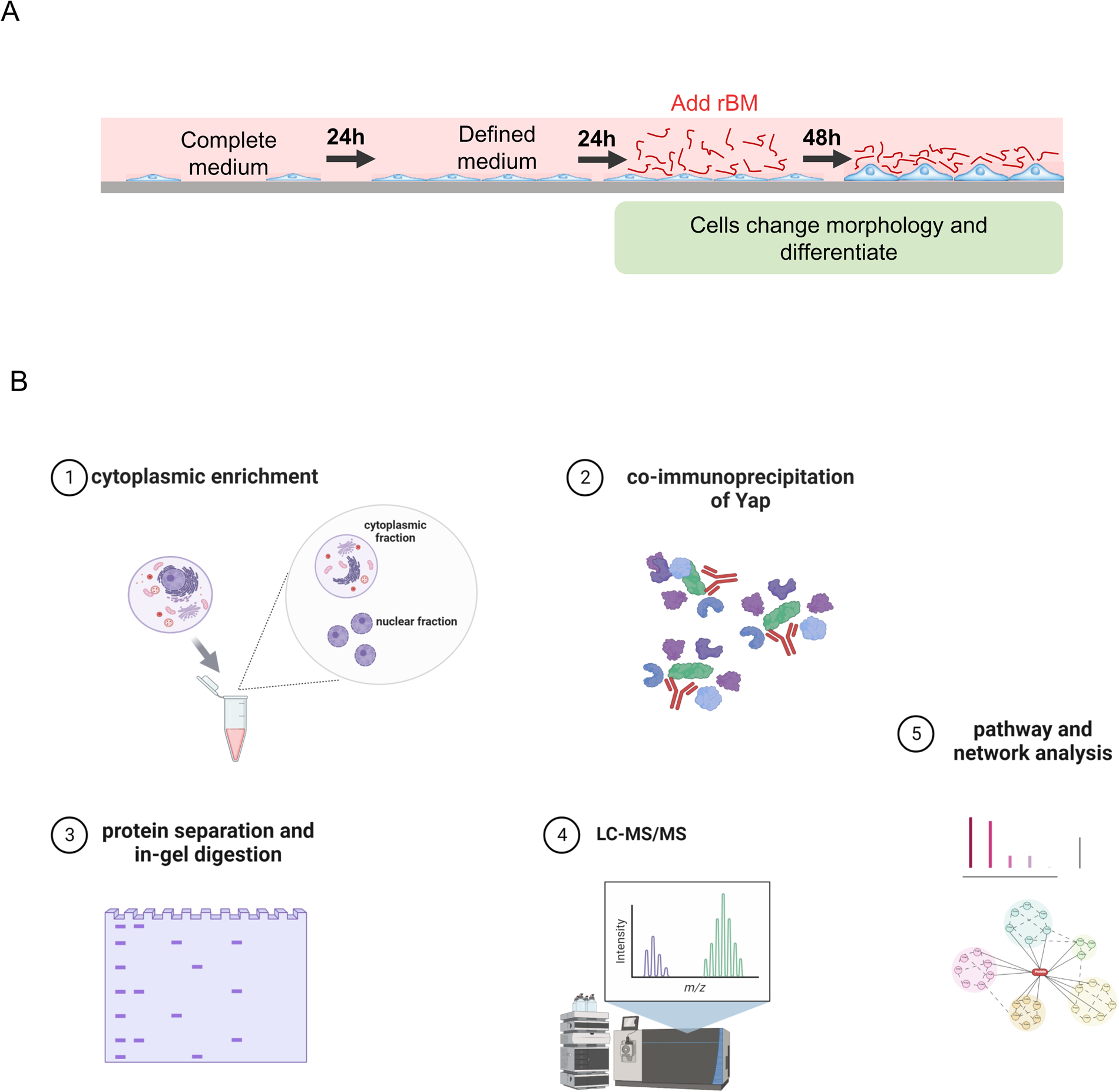
Schematics of experimental strategy to map YAP interactomes in the cytoplasm of mammary epithelial cells. A) Schematic of the rBM overlay assay where EpH4 cells are cultured until subconfluency and then treated with 2% rBM for 48 hours. B) Schematic of a workflow of Co-immunoprecipitation followed by proteomics to identify YAP interacting partners in the cytoplasm in cells treated or not with rBM.

For cell fractionation, we used a protocol with low stringency to yield a large number of interactants. The protocol resulted in appropriate enrichment of the cytoplasmic fraction, according to western blots for β-tubulin (a cytoplasmic protein) and lamin-B (a nuclear protein) (Suppl. Fig. 3A). To precipitate YAP, we used a monoclonal antibody and protein G covalently linked to Sepharose beads. An SDS-PAGE gel stained with silver nitrate revealed enrichment of a band at the expected molecular weight of YAP. (Suppl. Fig. 3B).

Next, we performed Co-IP of EpH4 cell-fractions treated or not with rBM, followed by mass spectrometry. We obtained 2716 hits from protein families (Suppl. Table 2). After excluding reverse peptides, common contaminants from mass spectrometry, and proteins identified in the precipitants of the non-specific IgG, 1565 hits remained, which were submitted to the online tool Venny to group unique proteins for each condition. We found a large number of hits that were exclusive to the different conditions, indicating that treatment with rBM results in differential YAP interactomes (Fig. 7A). On the other hand, proteins shared between the two treatments were not differentially abundant when their LFQ (label-free quantification) values were compared (Suppl. Fig. 4).

**Figure 7.**
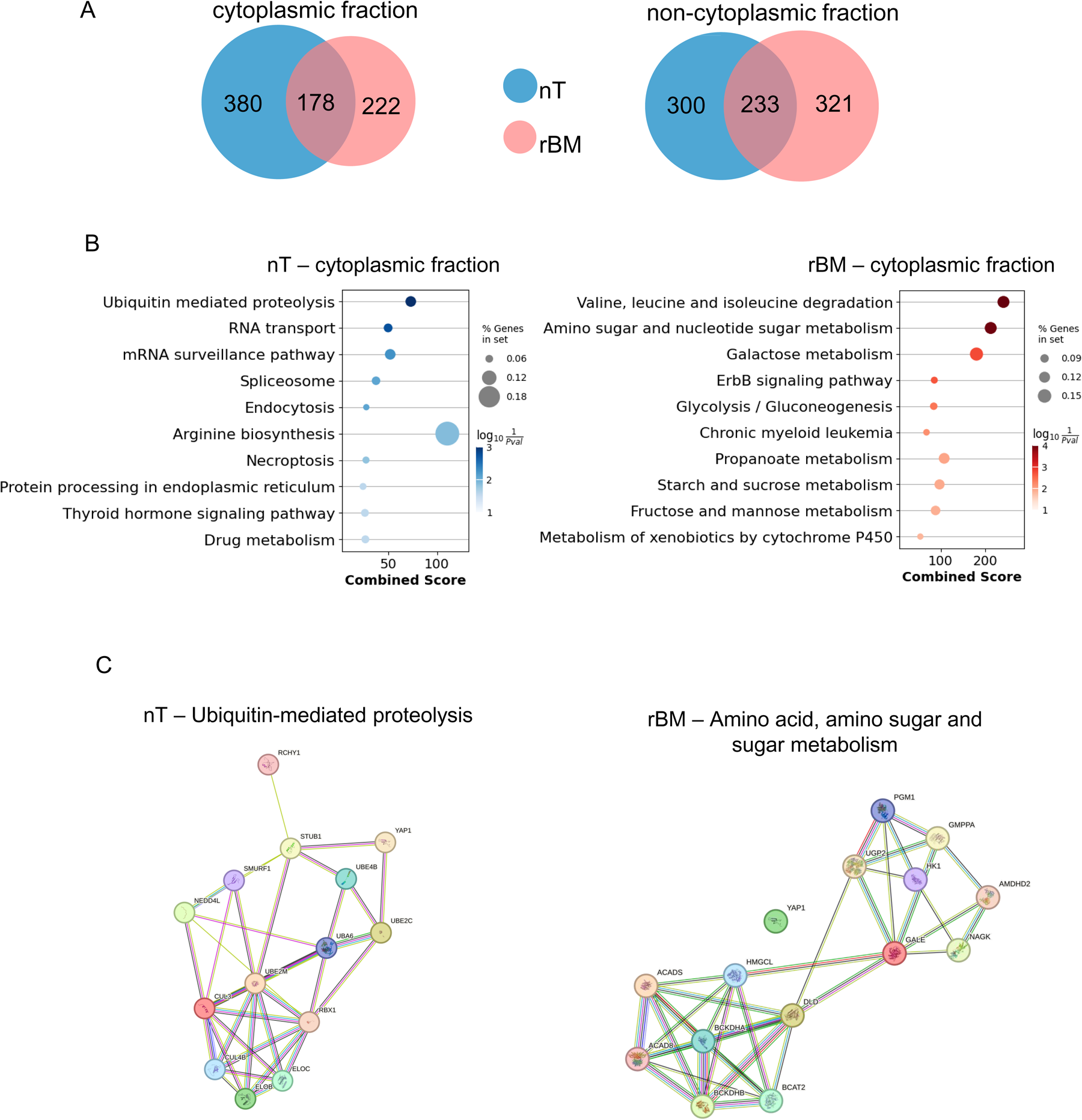
The basement membrane alters the interactome of YAP in the cytoplasm of mammary epithelial cells. A) Venn diagrams of YAP interactants in the cytoplasmic and non-cytoplasmic fractions of cells cultured in the presence or absence of rBM overlay. The data are shown as number of protein families. B) Cellular processes enriched in YAP interactomes in the cytoplasm of non-treated (nT) or rBM-treated epithelial cells. The analysis was performed with the Enrichr tool and using Kyoto Encyclopedia of Genes and Genomes (KEGG) pathways was used as the database. Only the interactants exclusively present in nT or rBM conditions were used for the analysis. The data in the bar graphs are shown as the cellular processes as a function of the -log_10_ of their enrichment p-value (color map). Dot sizes reflect the percentage of genes from the pathways found in each condition. C) String network analysis for top enriched pathways in nT and rBM-treated cells including YAP as a node. For the nT condition the average coefficient of local clustering was equal to 0.574, whereas for rBM-treated cells this parameter was 0.621. Both networks had a p-value of PPI enrichment < 1.0x10∧-16.

Using the enrichR tool (Kuleshov et al., 2016), we assessed the cellular processes in which unique proteins found in each condition may be involved (Fig. 6B, Suppl Tables 3-6). Ubiquitin-mediated proteolysis was the cellular process with the higher statistical significance in the cytoplasmic fraction of untreated cells, which agrees with the notion that cytosolic YAP is targeted for proteasomal degradation in subconfluent/growing cells (Kim & Jho, 2018; Zhao, Li, Tumaneng, et al., 2010).Unexpectedly, most proteins interacting with YAP in the cytosol of rBM-treated cells are involved in metabolic pathway processes, especially those related to amino acid and carbohydrate metabolism (Fig. 7B). Purine metabolism and pyrimidine metabolism were other identified cellular processes whose relationship to YAP has been reported (Santinon et al., 2018), but evidence about protein interaction hitherto had not been shown. For what we termed non-cytoplasmic fraction, we found the presence of terms related to the cell nucleus, plasma membrane, mitochondria, and other membranous organelles (Suppl. Fig. 4B).

Using STRING, a database of protein-protein interactions (PPI), we established interaction networks for the main cellular processes enriched in the cytoplasm (Fig. 7C). The 13 YAP interactants in the cytoplasm of non-treated cells and found in Ubiquitin-mediated proteolysis cellular term resulted in a PPI network with an average local clustering coefficient equal to 0.574 and a p-value PPI enrichment < 1.0x10^-16^, showing that the listed proteins have more interactions with each other than it would be expected from a protein list with similar size randomly generated from the genome, and indicating that the proteins are biologically connected as a group. However, only Stub1 and Ube2c had a previously reported association with YAP.

For cells treated with rBM, we combined the 14 proteins from the two top terms consisting of proteins involved in amino acid, amino sugar, sugar, and nucleotide metabolism that immunoprecipitated with YAP in the cytoplasmic fraction. The search in the STRING database also identified a network of well-documented interactions between the listed proteins, with an average coefficient of local clustering equal to 0.621 and a *p-value* of PPI enrichment < 1.0x10^-16^. Nevertheless, no documented YAP interactions were found with these metabolic proteins (Fig. 7C). The STRING platform is robust, but the interaction networks it builds are based only on evidence of interactions/associations from the literature. Because our strategy of mapping YAP interactants in the cytoplasm of cells in the context of the BM is novel, it is expected that the putative interactions we found have not been described. Therefore, it will be crucial in the future to design experiments to validate and explore the biological relevance of these molecular interactions.

## Discussion

Employing a comprehensive approach involving 2D and 3D cell culture, microscopy, bioimage analysis, proteomics, and bioinformatics, we investigated how the extracellular matrix (ECM) might influence the subcellular localization, functional behavior, and molecular associations of YAP, a critical element of the Hippo signaling pathway, in mammary epithelial cells.

We found that, as opposed to cells growing on 2D surfaces, YAP was predominantly cytoplasmic in non-malignant cells cultured in 3D-rBM and in the ducts of developing mammary epithelia, without noticeable changes in the protein levels in comparison to untreated cells and cells within the terminal end buds (TEB). It has been shown, in fact, that YAP can be either degraded or be bound to proteins associated with cell junction and ECM adhesion complexes depending in contexts of low mechanical stress (Gumbiner & Kim, 2014). But the cytoplasmic presence of YAP is interpreted as a strategy to block YAP translocation to the nucleus, where it would exert its transcriptional-related roles.

We hypothesized that YAP would have functions in the cytoplasm, and to generate insights to strength this hypothesis, we performed proteomics to identify YAP interactants in mammary epithelial cells exposed to an rBM. YAP interacted with different sets of proteins in the cytoplasmic compartments in mammary epithelial cells treated with rBM; YAP partners were several enzymes of metabolic pathways, especially amino acid, amino sugar and carbohydrate pathways. The mammary gland undergoes dramatic metabolic rewiring during development and especially in lactogenesis to sustain the high energetics demands of milk production (Hannan et al., 2023). This suggests that YAP not only is involved in nuclear and proliferative-related processes, but also acts as a modulator of metabolic changes. To the best of our knowledge, it remains elusive if YAP has non-nuclear nuclear functions and would act as a regulator of metabolic enzymes. After protein-protein interaction validations using biochemical and high-resolution imaging approaches, this open question could be addressed in future studies based on the molecular candidates our study unveiled.

In contrast to TEB cells, YAP was more cytoplasmic in the relatively quiescent ducts of the developing mammary gland, agreeing with what we observed in the 3D-rBM structures and cells submitted to ECM overlay. Moreover, TEBs showed a gene expression profile in which YAP-induced genes are overrepresented and enriched in comparison to the mature ducts, whereas a YAP-repressed gene set was only enriched and overrepresented in ducts. On the other hand, members of the Hippo pathway did not show significant changes in the TEB in duct transcriptomes. This body of evidence goes along with YAP localization pattern in the developing mammary gland, since nuclear YAP localization correlates with its transcriptional activity and active expression of YAP targets.

The pattern of YAP localization in TEBs that we unveiled is different from that previously reported (Chen et al., 2014), where YAP was detected in cap cells and body cells in 6-week TEBs. Possibly, the differences in the results between ours and their work was due to distinct techniques to stain YAP and the mouse strain used in the studies – while our work used immunofluorescence and BALB/C, theirs used immunohistochemistry (IHC) and C57BL/6 background. Another important factor is that their work did not quantify the IHC YAP staining. Nevertheless, this important article showed that YAP was not essential for mammary gland morphogenesis and cancer initiation, but it was crucial for lactogenesis and tumor progression, underscoring the complex temporal and contextual regulation of this Hippo pathway effect in the biology of the mammary gland.

Upon increased mechanical stress, YAP and TAZ translocate into the nucleus and associate with TEAD transcription factors to mediate targeted gene transcription (Cai et al., 2021).. However, in distinct mechanical contexts YAP/TAZ nuclear localization and activation do not seem to rely solely on the Hippo pathway. Rather, regulation of YAP/TAZ depends on the tension built in the actomyosin cytoskeleton (Das et al., 2016; Dupont et al., 2011). In fact, the BM induces changes in cell morphology by altering the cytoskeleton (Fiore et al., 2017; Glukhova & Streuli, 2013; Streuli et al., 1995). Thus, it is possible that the actin cytoskeleton may relay the BM signals that influence YAP localization. In contact with the BM, normal epithelial cells display a program of tissue morphogenesis that culminates with the assembly of quiescent acini. However, malignant cells form proliferative and tumor-like structures and possess higher number of stress cytoskeleton fibers and do not polarize with a clear apical domain (Rizki et al., 2008). This differential response to rBM in respect to cytoskeleton organization and polarization along with evidence that the cytoskeleton regulates YAP/TAZ localization and function might explain why malignant cells do not show the same subcellular localization for YAP as their nonmalignant counterparts.

Also importantly, although we have not seen striking changes in the mRNA levels of key Hippo components in the developing mammary gland and in protein and in protein phosphorylation levels of non-malignant cells exposed to an rBM, we can not rule out the involvement of Hippo in YAP localization regulated by the BM. Furthermore, we observed a discrepancy between mRNA levels of MST2 and YAP with their protein levels in 3D-cell cultures. This indicates that at the time point we analyzed, at least for mRNA of these Hippo components, our assay does not reproduce what was seen for the developing mammary gland and these genes might differentially regulated by the BM in specific contexts. This should be addressed in future studies including genetic and pharmacological perturbation of the Hippo pathway. Likewise, the intriguing finding that in the presence of rBM, YAP not only is predominantly retained in the cytoplasm, but it is also localized in the nuclear periphery raise the question if in growth suppressive contexts, YAP would have specific transcriptional actions in genes localized in proximity to the nuclear envelope.

The Hippo pathway and YAP have been implicated in the development of diseases, particularly cancer and conditions of disrupted tissue regeneration, and one of the hallmarks of adenocarcinomas is the disruption of the basement membrane along with loss of epithelia polarity and proliferative quiescence (Cerqueira et al., 2022; Fiore et al., 2017; Tomasin et al., 2023; Zhao, Li, Lei, et al., 2010). Our findings that human breast malignant cells (T4-2 cells) in 3-rBM presents elevated concentrations of nuclear YAP suggests that the highly proliferative capacity of malignant cells, even in a growth-suppressive environment like that provided by basement membrane components, might be linked with deregulated YAP, both in the nucleus and in the cytoplasm. These mechanisms should be further addressed and leveraged to pave the way for novel strategies to fight cancer.

## Material & Methods

### Cell lines and culture conditions

EpH4 cells (murine mammary non-malignant epithelial cell line) were cultured in DMEM-F12 (Gibco; cat. #12500-039) medium supplemented with 2% heat-inactivated fetal bovine serum (Gibco; cat. #12657-029), 5 μg/mL insulin from bovine pancreas (Sigma; cat. #I5500), and 50 μg/mL gentamicin (Gibco; cat. #15710-064). Cells from the HMT-3522 series (nonmalignant S1 and tumoral T4-2 cells) were cultured as previously described (Fiore et al., 2017) in DMEM/F-12 (Gibco; cat. #11330032) supplemented with 25 ng/mL insulin, 10 μg/mL human apo-Transferrin (Sigma-Aldrich; cat. #T1147), 2.6 ng/mL sodium selenite (Sigma-Aldrich; cat. #S-5261), 0.1 nM β-estradiol (Sigma-Aldrich; cat. #E2758), 1.4 μM hydrocortisone (Sigma-Aldrich; cat. #H4001), 5 μg/mL prolactin from sheep pituitary (Sigma; cat. #L6520), and 10 ng/mL epidermal growth factor from murine submaxillary gland (Sigma-Aldrich; cat. #E4127; added exclusively to S1 cells). All cell lines used in this work were a kind gift from Dr. Mina J. Bissell, from Lawrence Berkeley Laboratory (Berkeley, CA, USA) and were routinely tested for mycoplasma contamination.

For “on top” 3D cell culture experiments, 200 µL of ice-cold reconstituted basement membrane (rBM) (growth-factor reduced Matrigel, Corning cat; #354230) was distributed in 24-well plates and incubated at 37°C for gelification. Five thousand S1 and T4-2 cells were resuspended in a defined medium containing 5% rBM and seeded over the rBM coating. Phenotypical reversal of T4-2 cells was performed by adding 10 µM Tyrphostin AG 1478 (Sigma, cat; # 175178-82-2) to the medium on day zero of 3D culture as previously described (Fiore et al., 2017; Wang et al., 2002). Cells were cultured for 5 days, until 3D structure assembly.

For 3D cell culture on non-adhesive plates, EpH4 cells were seeded (10000 cells/cm^2^) in a defined medium (DMEM/F-12 supplemented with 5 μg/mL insulin from bovine pancreas, 1 μg/mL hydrocortisone, and 50 μg/mL gentamicin) on a plate precoated with 0.8 mg/cm^2^ of Poly(2-hydroxyethyl-methacrylate) (PolyHEMA; Sigma-Aldrich cat. #P3932). Cells were cultured for 48h, collected by centrifugation, and replated in a differentiation medium (the same defined medium previously used supplemented with 3 µg/mL prolactin from sheep pituitary and 2% rBM) on a PolyHEMA-coated plate. After 48-72 hours acini were collected for further analysis.

For EpH4 ECM-overlay experiments, cells were seeded in growth medium at a density of 2x10³/cm². After 24 hours, the medium was replaced with a defined medium (same as used in 3D experiments). Cells were cultured for another 24 hours, when the medium was replaced by the differentiation medium, and cells were cultured for 48 hours.

### Bioinformatics analysis of the developing mammary gland transcriptomics

The data for the gene expression analysis of mouse mammary TEB versus duct (Kouros-Mehr & Werb, 2006), GSE2988, were downloaded from the Gene Expression Omnibus database (https://www.ncbi.nlm.nih.gov/geo/). Genes with M value > 0.6 (1.5-fold change) and adjusted P value < 0.05 (Benjamini–Hochberg correction) were considered differentially expressed in TEB or mature ducts relative to distal stroma (Benjamini,1995). TEB ratio = M(TEB) – M(duct), converted to fold change. Duct ratio = M(duct) – M(TEB), converted to fold change. The EnrichR tool (Kuleshov et al., 2016) (https://maayanlab.cloud/Enrichr/) was used for overrepresentation analysis (ORA). The lists for YAP-regulated genes were obtained from the literature: induced by YAP in 3T3_Zhao (Zhao et al., 2007), induced by YAP_Zhang (Zhang et al., 2009), YAP_Dupont (Dupont et al., 2011), YAP conserved signature_Cordenonsi (Cordenonsi et al., 2011) and repressed by YAP in 3T3_Zhao (Zhao et al., 2008). Gene set enrichment analysis (GSEA) was performed with the GSEA software (https://www.gsea-msigdb.org/gsea/index.jsp). The heatmap plots to assess the gene expression variation of the Hippo pathway genes and genes induced or repressed by YAP were generated with the online tool Morpheus (https://software.broadinstitute.org/morpheus/).

### Cell fractionation and co-immunoprecipitation

Cells were detached from the plate with a cell scraper and washed twice with cold PBS. For fractionation, cells were resuspended in PBS containing protease/phosphatase inhibitors and subjected to 3 cycles of quick freezing in liquid nitrogen followed by thawing on ice and trituration by pipetting. The lysate obtained was centrifuged at 13000xg for 20 min at 4 °C and supernatant was collected as cytoplasmic enriched fraction. The remaining pellet was washed once with PBS, resuspended in PBS + 0.5% Nonidet-P40 and protease/phosphatase inhibitors, and sonicated on ice (5 seconds of sonication at the amplitude of 20 followed by 30 minutes of rest on ice). This lysate was centrifuged at 15,000xg for 20 minutes at 4°C and supernatant was collected as the cytoplasmic-depleted fraction (or non-cytoplasmic fraction).

For immunoprecipitation, 350 μg of each fraction was cleared by incubation with 10 μL of Protein G coupled to agarose beads pre-washed and equilibrated with PBS + 0.1% Nonidet-P40 for 2 hours at 4°C under agitation. After clearance, the remaining supernatant was incubated overnight with 20 μg of a specific monoclonal antibody for YAP (Santa Cruz, cat. #sc-101199) and 10 μL Protein G coupled to agarose beads and (Santa Cruz Biotechnology). Then beads were washed with ice cold PBS three times and the final resuspension was done in Laemmli Buffer (50mM Tris-HCl, pH 6.8, 2%SDS, 10% glycerol, and 100mM β-mercaptoethanol) for gel electrophoresis followed by western blotting or sample preparation for mass spectrometry.

### Reverse transcription quantitative PCR (RT-qPCR)

Total RNA was extracted from cells using TRIzol Reagent (Invitrogen). Reverse transcription of total RNA was done using Superscript II kit (Invitrogen) and both random hexamers (Invitrogen) and oligo d(T)20 (Invitrogen) sequences. qPCR analysis was performed using Power SYBR Green PCR Master Mix Reagent (Invitrogen) in a thermocycler 7500 Real-Time PCR System (Applied Biosystems). Ct values were normalized to reference gene *B2m*. Designed primers: *Ankrd1* (Fw-GAGACACCCCACTGCATGAT; Rv-TTCCCAGCACAGTTCTTGACC), *Yap* (Fw-TTCGGCAGGCAATACGGAAT; Rv-CATCCTGCTCCAGTGTAGGC), *Stk3* (Fw-GAGAAGCTTGGAGAAGGGTCTT; Rv-GCAACCACTTGACCAGATTCC), *Ect2* (Fw- GCACATCATCCTTAGCAGGTAT; Rv-GTCAGCGTCTTGTTGAAGCAT), *Csn2* (Fw-CATTTACTGTATCCTCTGAGACTGA; Rv-TGTGACTGGATGCTGGAGTG) *B2m* (Fw-CACTGAATTCACCCCCACTGA; Rv-TGTCTCGATCCCAGTAGACGG).

### Western Blotting

EpH4 cell structures were collected by centrifugation, washed twice with cold PBS and lysed for 30 minutes on ice with RIPA buffer (50mM Tris-HCl, pH 8, 150mM NaCl, 0.1% SDS, 1% sodium deoxycholate, 1% Nonidet P-40, and 0.5mM EDTA) containing protease (Roche; cat. #) and phosphatase (Sigma-Aldrich; cat. #) inhibitors. After centrifugation at 15,000xg, 4°C, for 20 minutes, supernatant was collected, and its protein content was quantified with DC Protein Assay kit (Bio-Rad; cat. #5000112). Fifteen μg of cell lysate were loaded into 10% polyacrylamide gels for protein electrophoresis and then blotted to PVDF membranes. Membranes were blocked with 5% BSA diluted in TBS-T buffer (50mM Tris-HCl, pH 7.4, 150mM NaCl, 0.1% Tween-20) for 1 hour, washed with TBS-T (3 washes, 10 minutes each) and incubated overnight with primary antibodies at 4°C. Membranes were washed as previously and then incubated with the appropriate HRP-conjugated secondary antibody for 1 hour at room temperature. The chemiluminescence signal was detected with Chemidoc Imaging System (Bio-Rad) and using the SuperSignal West Dura substrate (Thermo), according to manufacturer’s instructions. The detected bands were quantified with the software ImageLab 6.0 (Bio-Rad).

### List of antibodies used in this work

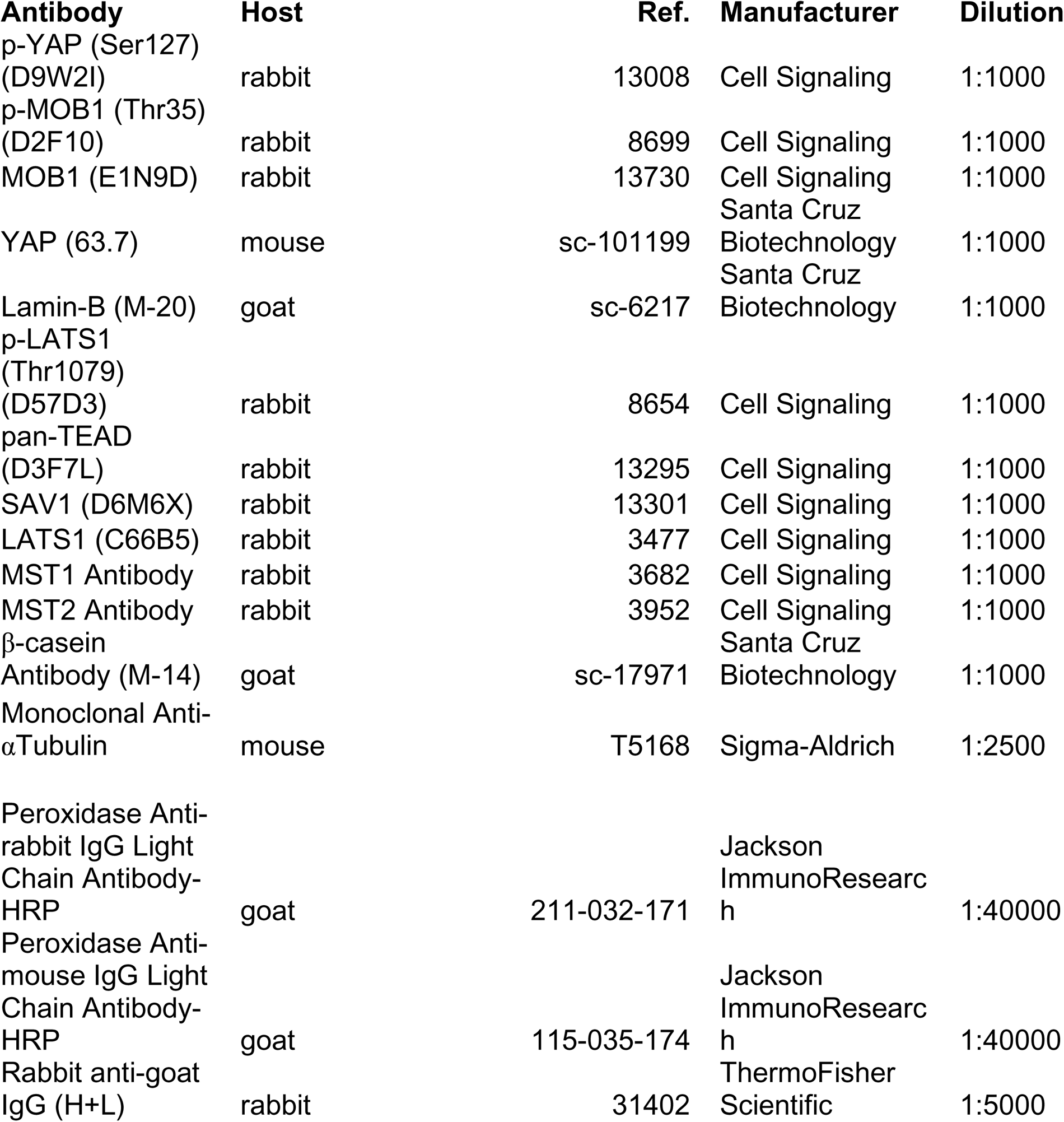
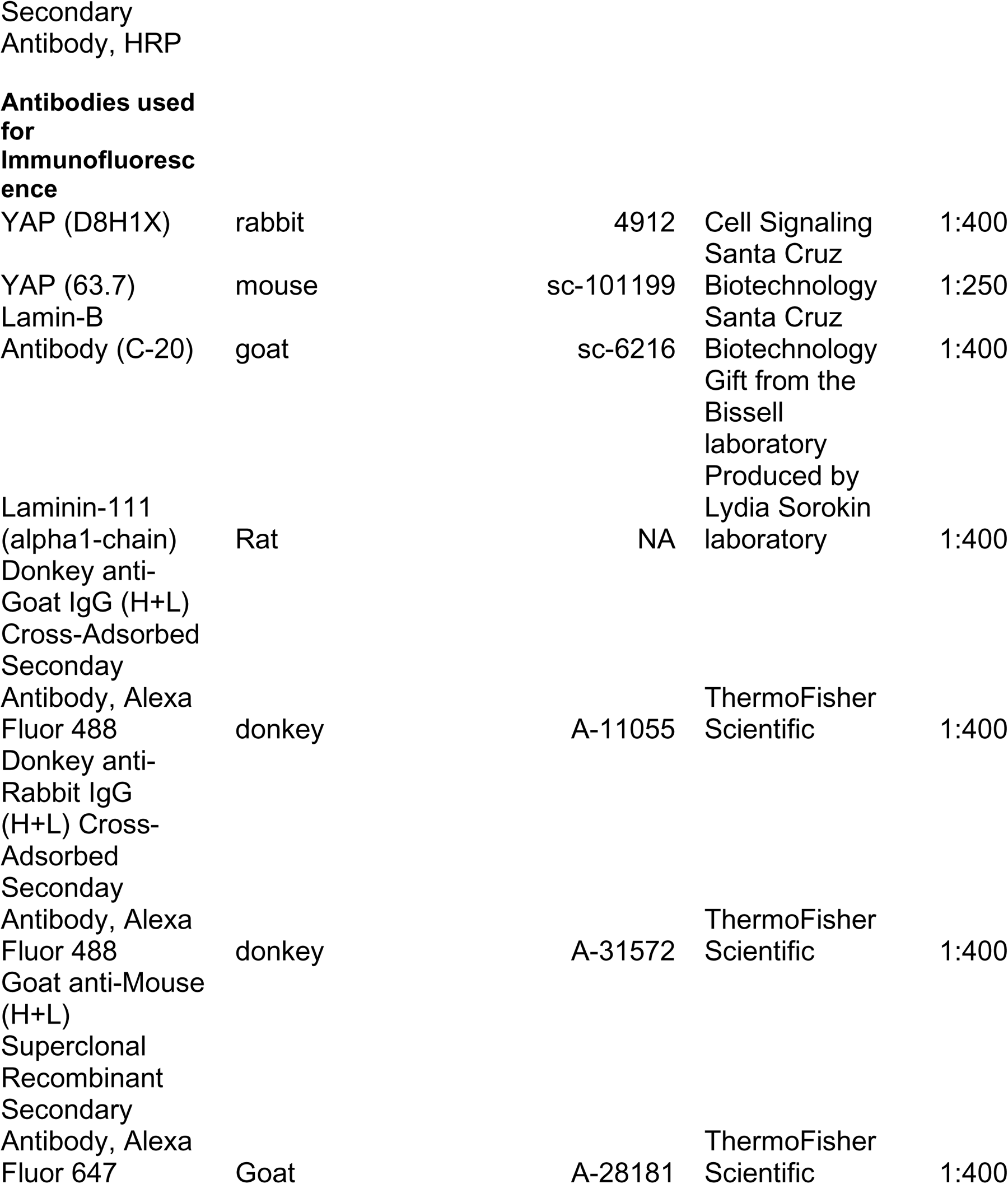

### Immunofluorescence and image analysis

Cells and 3D structures were fixed with 4% paraformaldehyde (PFA) diluted in PBS pH 7.4 for 10 minutes for cells on 2D or 30 minutes for 3D structures. Residual PFA reactivity was quenched with PBS-10 mM glycine and cells were permeabilized with 0.5% Triton X-100 in PBS (10 minutes for 2D cells and 30 minutes for 3D structures). Cells were incubated in blocking solution (3% Bovine Serum Albumin (BSA) with 5% goat serum in PBS) for 1 hour at room temperature, followed by overnight primary antibody incubation in blocking buffer. After 3 washes with PBS (5 minutes each), the samples were incubated with the appropriate secondary antibody (Alexa Fluor-488, Alexa Fluor-594, Alexa Fluor-647 at concentration 1:400), diluted in 3% BSA in PBS for 45 minutes at room temperature in the dark.

For immunofluorescence on tissue sections, 5-week BALB/C female mice were euthanized and had their 4^th^ mammary glands collected. The experimental procedures involved in this work were approved by the Ethics Committee for Animal Welfare from the Institute of Chemistry/USP (protocol #16/2015). The glands were snap frozen in Tissue Tek Optimal Cutting Temperature, sectioned with a cryostat, and submitted to immunofluorescence for detection of YAP and laminin-111. Sections were fixed with 4% PFA in PBS. The residual PFA reactivity was quenched with PBS–25 mM glycine, and the tissue sections were permeabilized with 0.5% Triton X-100 in PBS for 15 minutes, incubated in a blocking solution (1% BSA with 5% goat serum) in PBS for 1 hour at room temperature, followed by overnight incubation with primary antibodies. The tissues were washed with PBS and incubated with the appropriate secondary antibody conjugated to Alexa Fluor 488 or Alexa Fluor 594, diluted in 1% BSA in PBS, for 45 minutes at room temperature in the dark.

The primary antibodies used in this work for immunofluorescence were YAP (Cell Signaling, cat #4912S, 1:400), YAP (Santa Cruz cat #sc-101199, 1:250), Lamin-B (Santa Cruz, cat #sc-6217). and, Laminin-111 1:400 (anti-laminin α1 chain - clones 198 and 200 produced by Lydia Sorokin). For all samples, the nucleus was counterstained with 0.5 µg/mL 4’6-diamidino-2-phenylindole (DAPI) for 10 minutes and slides were mounted with Prolong® Gold (Life Technologies).

For quantification of YAP in glandular structures (TEB and duct) images were acquired in a microscope Zeiss LSM (Laser Scanning Microscope) using a 40x/0.75NA objective. For quantification, we defined the mammary gland epithelia containing TEB in 3 zones: Zone A: 40µm from the most distal region of the TEB, Zone B as 40 µm after the proximal limit of Zone A, and Zone C (includes duct) as the next 80 µm. To quantify YAP intensity, we used Image J software (http://imagej.nih.gov/ij/). In brief, for each image the channels were separated and the images with DAPI staining were binarized and segmented to generate nuclear masks. The mean fluorescence intensity of YAP was quantified in each nuclear mask. To quantify YAP in the cytoplasm, we increased the contour of the nuclear masks concentrically by 0.7 µm and measure the fluorescence intensity in this new selected area that we termed total area (area that encompasses the nucleus and part of the cytoplasm). Then, the mean value of intensity for the cytoplasm was defined by (intensity total - nuclear intensity / total area - nuclear area) and the nucleus cytoplasm ratio (nuclear YAP intensity/cytoplasmic YAP intensity) was determined.

Images for quantification of YAP in EpH4 cells in 2D and 3D, cultured in the presence or absence of rBM, were captured in a widefield Leica DMi8 microscope using a 63x/1.4NA oil objective and deconvoluted in Leica’s LASX software. Images for quantification of YAP in 3D structures of S1 and T4-2 cells were captured in a Zeiss LSM (Laser Scanning Microscope) microscope using a 40x/0.75NA objective. To quantify nuclear YAP intensity in both 2D cells and 3D structures, we used the same strategy described above for YAP quantification in TEB/duct.

### Phase contrast and morphological quantification of 3D structures

To measure the acinogenesis process of EpH4 3D structures, we used phase contrast microscopy. Images were captured with a DMi1 microscope (Leica Microsystems), with a camera attached to the microscope, using LAX software (Leica Microsystems). The number of structures with acinar morphology (presence or absence of lumen), the size of the structures (calculated through expanded area), and the circularity of the structures were analyzed. Circularity (C) was determined by the formula:

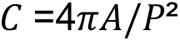

Where C represents circularity value, A stands for the area of the median plane, and P represents the perimeter. The analyses were conducted using ImageJ2 v1.52f software provided in the Fiji distribution (Schindelin et al., 2012), along with the assistance of PHANTAST v0.2 plugin (Jaccard et al., 2014). In summary, after calibration to the micrometric scale, the images were converted to 32-bit format, and structure delineation was performed using the PHANTAST plugin (parameters: sigma = 1.20; epsilon = 0.25; halo correction = enabled; selection overlay on original image = enabled; new mask image = enabled). The generated mask was corrected with the fill holes tool and analyzed using the analyze particles tool (parameters: size = 500-infinity; area = enabled; shape descriptors = enabled; perimeter = enabled). The presence of lumens in the structures was determined through visual inspection of phase contrast photomicrographs, and the counting was done manually.

### Sample preparation, mass spectrometry and data analysis

Samples from Co-IP experiments were resolved by SDS-PAGE in a gradient polyacrylamide gel Bolt™ 4-20% Bis-Tris Plus (Invitrogen). After running, gels were stained with Colloidal Coomassie R-250 for lane demarcation. Each lane was sectioned into 4 pieces, and each piece was separately processed and analyzed by mass spectrometry. Gel fragments were incubated 3 times with 100 mM ammonium bicarbonate (Sigma-Aldrich; cat. #09830) to wash away excess SDS and change the buffer. Reduction of disulfide bonds and alkylation of cysteine residues was performed by incubating gel fragments in 10 mM 1,4-Dithio-D-threitol (GE Healthcare; cat. #17-1318-02) at 50°C for 1 hour followed by a light-protected incubation with 55 mM iodoacetamide (GE Healthcare; cat. #RPN6302V) at room temperature for 45 min. Proteins were digested in-gel overnight at 37°C with Sequencing Grade Modified Trypsin (Promega; cat. #V5111) diluted in 200uL of digestion buffer (40 mM ammonium bicarbonate, and 10% acetonitrile (Merck; cat. #1.00029). Gel fragments were dehydrated with pure acetonitrile for peptides extraction and the recovered supernatants were totally dried in a vacuum concentrator (SpeedVac). The recovered tryptic peptides were resuspended in 0.1% trifluoroacetic acid (Sigma-Aldrich; cat. #T6508) and desalted with ZipTips C18 Pipette Tips (Millipore; cat. #ZTC185096) according to manufacturer instructions. The eluted peptides were dried and stored at an -80°C freezer until analysis.

The purified peptides were analyzed using the reverse phase liquid chromatography system Easy-nLC 1200 UHPLC (Thermo Scientific) coupled to an Orbitrap Fusion Lumos Tribrid Mass Spectrometer (Thermo Scientific) equipped with a nanospray Flex NG ion source (Thermo Scientific). Samples were loaded onto a trap column (Acclaim PepMap 100, C18, 3um, 75um x 2cm, nanoViper; Thermo Scientific) with solvent A (0.1% formic acid) at 500bar, and then were eluted to a C18 column (Acclaim PepMap RSLC, C18, 2 µm, 75 µm x 15 cm, nanoViper; Thermo Scientific) at a flow rate of 300nL/min over 90min. Peptides were separated using a linear gradient of solvent of 5-95% solvent B (0.1% formic acid in acetonitrile). After peptide separation, the column was washed for 12 minutes with 100% solvent B and the system was re-equilibrated with 100% solvent A. The ion source was operated in positive ESI mode with capillary temperature set at 300°C, and S-Lens RF level at 30%. A full MS scan was followed by data dependent MS2 scans in a 3-second cycle time. Both MS and MS2 scans were performed in the Orbitrap analyzer. Precursor ions were fragmented by HCD with a normalized collision energy of 30. Full scan MS spectra (m/z 300-2000) of peptides were acquired at a resolution of 120000; m/z=445.12,003 was used as a lock mass. Instrumental conditions were checked between samples using 100 fmol of a tryptic digest of BSA as a standard. Samples vestiges were completely removed between samples.

Raw data was processed using MaxQuant suite. Raw file spectra were searched against the *Mus musculus* Uniprot reference database (reference UP000000589; organism ID 10090; downloaded in August, 2017). Trypsin was set as the digestion enzyme, with a maximum of 2 missed cleavages allowed. Cysteine carbamidomethylation was selected as a fixed modification and methionine oxidation and N-terminal acetylation were selected as variable modifications. MaxQuant default options were used for all the other parameters. All proteins identified with one or more peptides were selected for the subsequent analysis. After the exclusion of potential contaminants and reverse peptides, the shared and exclusive proteins for each condition was defined with the online tool Venny (http://bioinfogp.cnb.csic.es/tools/venny/). The EnrichR module of the python package GSEApy (https://gseapy.readthedocs.io/en/latest/introduction.html#) along with the Kyoto Encyclopedia of Genes and Genomes (KEEG; Mouse 2019) database was used for overrepresentation analysis (ORA). The interaction networks were done with the platform STRING v12 (Szklarczyk et al., 2015).

## Acknowledgements

The authors would like to thank Celia Ludio for technical assistance. We thank the Central Analitica - IQUSP for the access to equipment and for the assistance provided. The authors are also grateful to Graziella Eliza Ronsein and Giuseppe Palmisano for insightful inputs in the proteomics experimental design and data analysis.

## Data availability statement

Tables of the gene set signatures used in this work, the LFQ for all proteins found in the mass spectrometry analysis, the exclusive protein families found for each condition, and the Enrichr output were deposited at https://zenodo.org/records/10845652.

