## Supplementary figures and images for "The basement membrane regulates the cellular localization and the cytoplasmic interactome of Yes-Associated Protein (YAP) in mammary epithelial cells"

### suppl. figures

Suppl. Figure 1

A

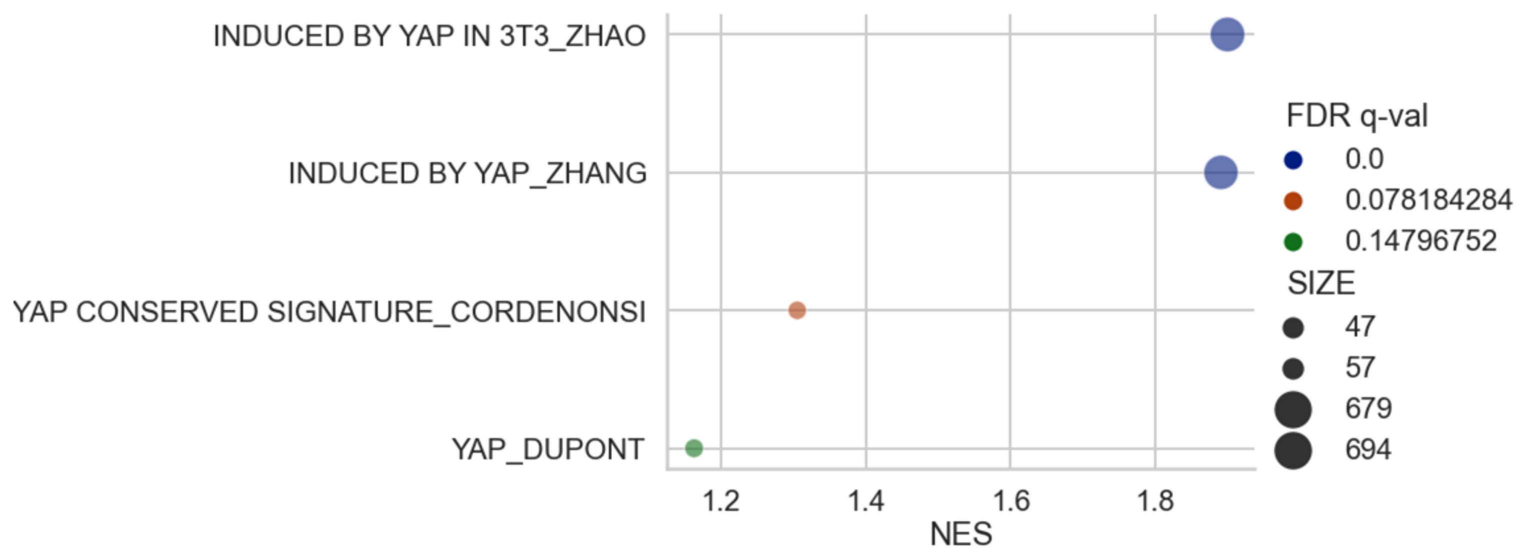


B

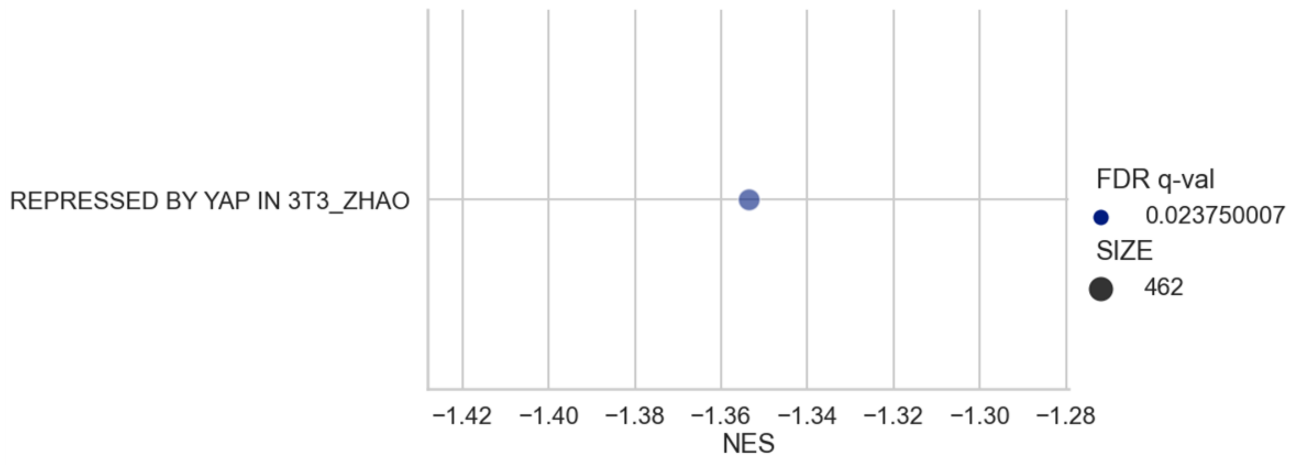

Suppl. Figure 2

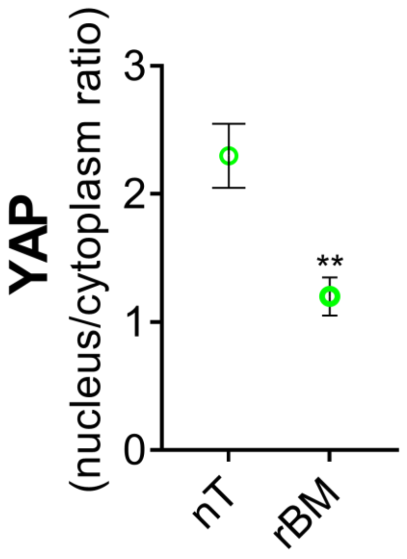


Suppl. Figure 3

A

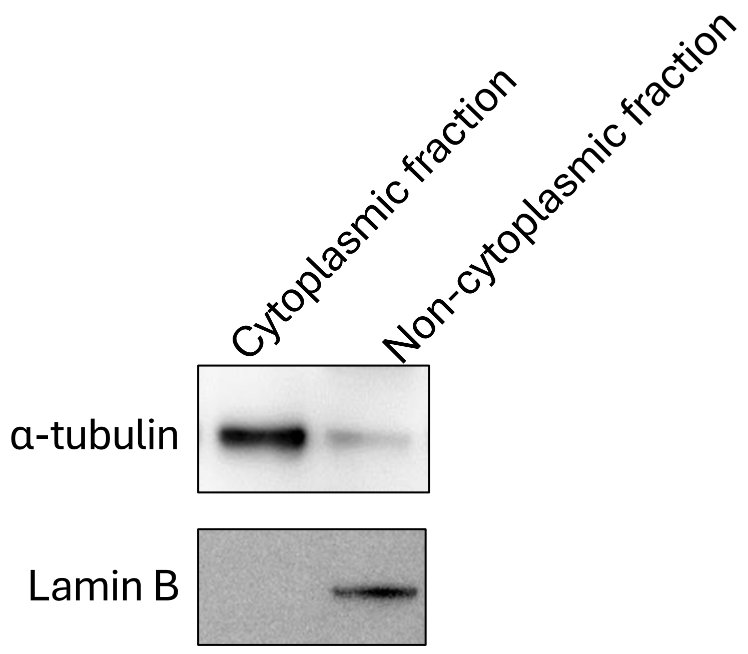


B

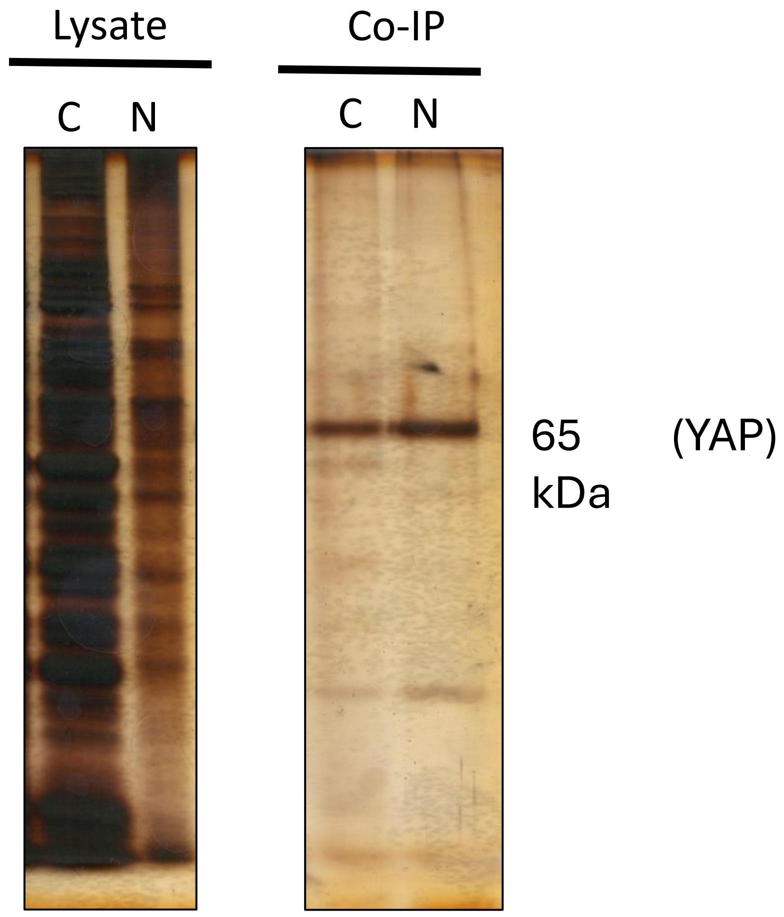

Suppl. Fig. 4

A

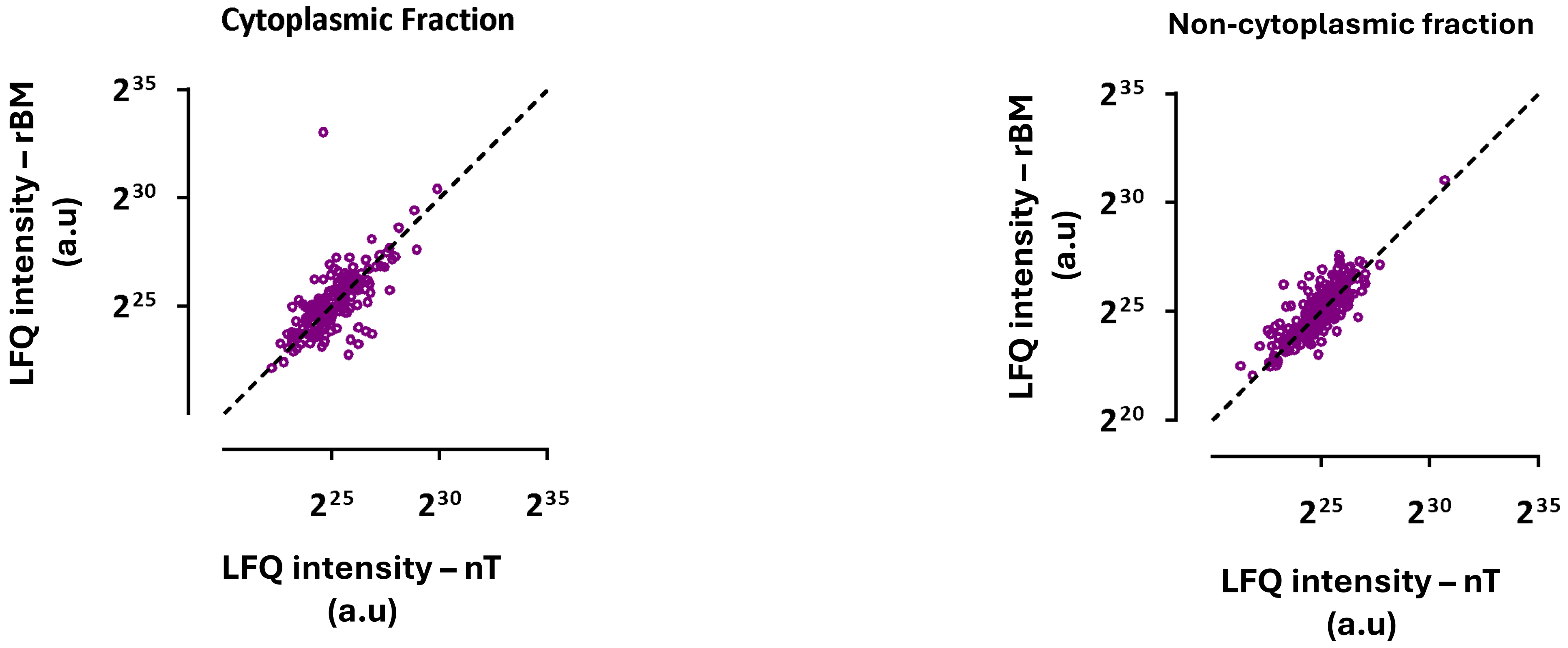


B

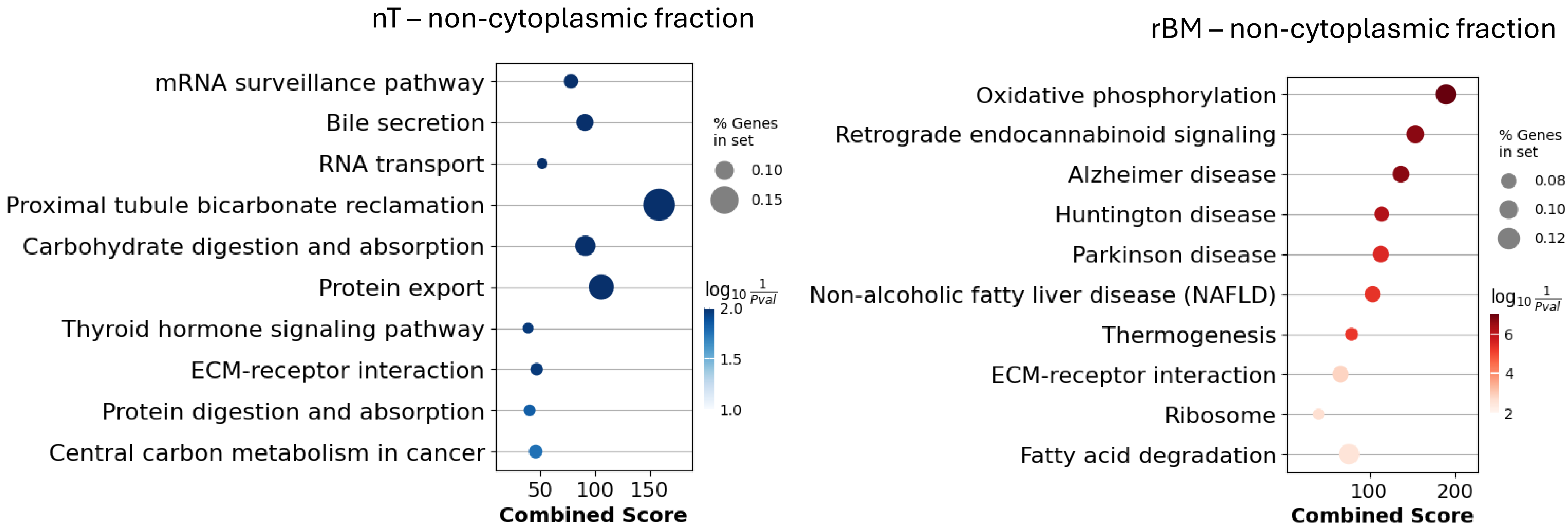
